# Logical modelling and analysis of cellular regulatory networks with GINsim 3.0

**DOI:** 10.1101/289298

**Authors:** Aurélien Naldi, Céline Hernandez, Wassim Abou-Jaoudé, Pedro T. Monteiro, Claudine Chaouiya, Denis Thieffry

**Affiliations:** Computational Systems Biology Team, Institut de biologie de l’École Normale Supérieure (IBENS), École Normale Supérieure, CNRS, INSERM, PSL Université, Paris, France; INESC-ID/Instituto Superior Técnico, University of Lisbon, Lisbon, Portugal; Instituto Gulbenkian de Ciência, Oeiras, Portugal

**Keywords:** Regulatory network, logical model, discrete dynamics, regulatory circuit, p53-Mdm2 network

## Abstract

The logical formalism is well adapted to model large cellular networks, for which detailed kinetic data are scarce. This tutorial focuses on this well-established qualitative framework. Relying on GINsim (release 3.0), a software implementing this formalism, we guide the reader step by step towards the definition, the analysis and the simulation of a four-node model of the mammalian p53-Mdm2 network.

## 1 Introduction

The logical formalism is becoming increasingly popular to model cellular networks [3, 31]. Here, we focus on the framework developed by René Thomas and colleagues, which includes the use of multilevel variables when functionally justified, along with sophisticated logical rules or parameters [46, 47].

This approach has been applied to the study of a wide range of networks controlling, for example, the lysis-lysogeny decision of the bacteriophage *λ* [45], the specification of flower organs in arabidopsis [4, 29], the segmentation of drosophila embryo [27, 39, 40, 41], the formation of compartment in drosophila imaginal disks [21, 22], drosophila egg shell patterning [18], cell cycle in mammals and yeast [16, 17, 48], the specification of immune cells from common progenitors [12, 28], the differentiation of T-helper lymphocytes [1, 25, 30], neural differentiation [14], as well as cancer cell fate decisions [8, 19, 23, 37, 38], etc.

In order to ease access to logical modelling by biologists, this protocol proposes a stepwise introduction to the framework, relying on its implementation into the software GINsim (release 3.0). The following section introduces the biological system used as an illustration. Next, in Section 3, we proceed with the stepwise construction and analysis of a logical model. The article then ends with some conclusions and prospects.

## 2 The p53-Mdm2 network

The transcription factor p53 plays an essential role in the control of cell proliferation in mammals by regulating a large number of genes involved notably in growth arrest, DNA repair, or apoptosis [49]. Its level is tightly regulated by the ubiquitin ligase Mdm2. More precisely, nuclear Mdm2 down-regulates the level of active p53, both by accelerating p53 degradation through ubiquitination [7] and by blocking the transcriptional activity of p53 [15, 34]. In turn, p53 activates Mdm2 transcription [5] and down-regulates the level of nuclear Mdm2 by inhibiting Mdm2 nuclear translocation through inactivation of the kinase Akt [26]. Finally, high levels of p53 promote damage repair by inducing the synthesis of DNA repair proteins [20].

Given its key role in DNA repair and cell fate control, various groups have modelled this network by using different formalisms, including ordinary differential equations [11, 50], stochastic models [35, 36, 43], hybrid deterministic and stochastic models [24], as well as logical models [2, 10].

In this protocol, we rely on a refined version of a logical model presented by Abou-Jaoudé et al. [2], involving the protein p53; the ubiquitin ligase Mdm2 in the cytoplasm; the ubiquitin ligase Mdm2 in the nucleus; and DNA damage (see Figure 1).

**Figure 1:**
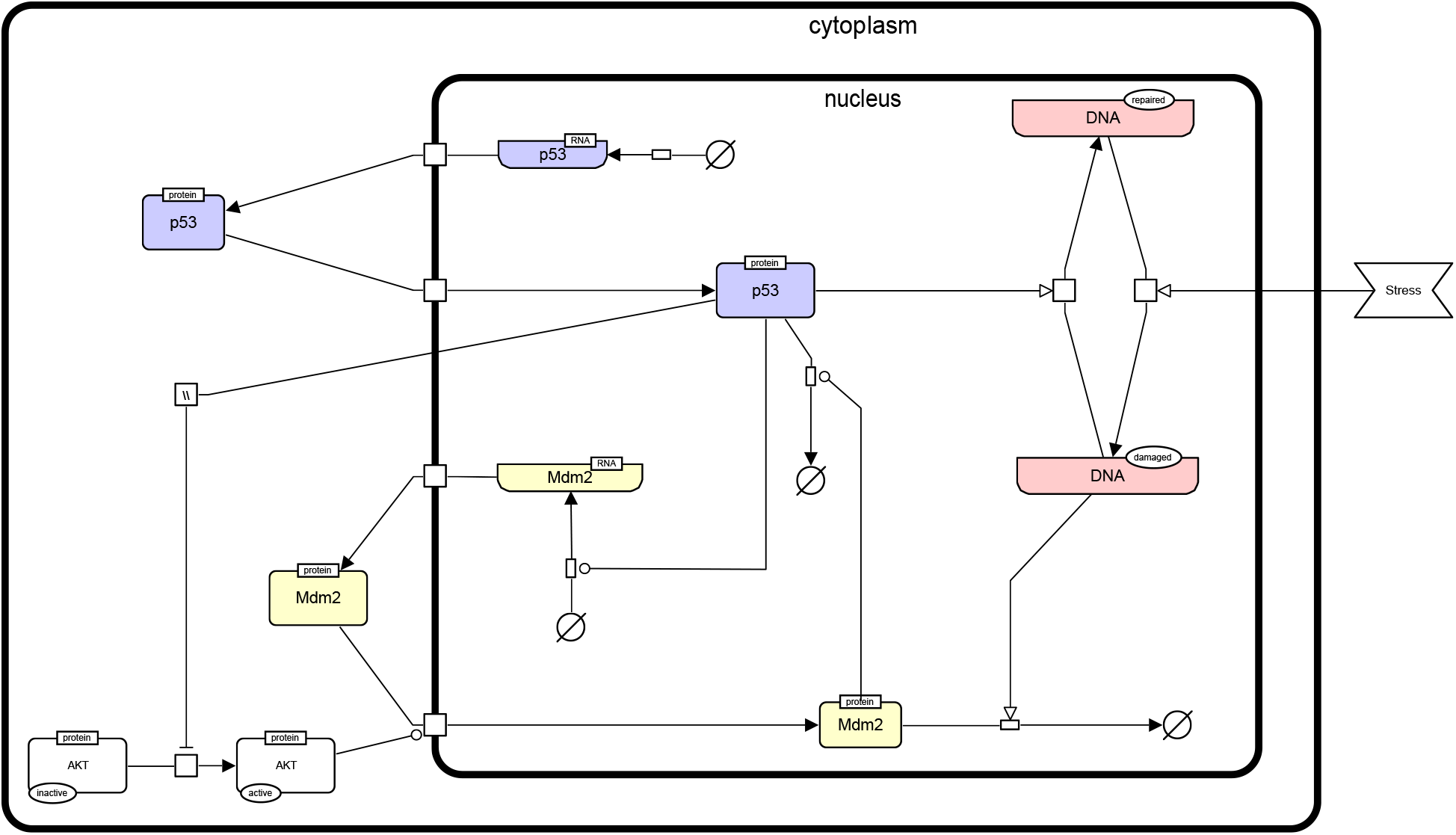
The p53-Mdm2 network. This figure describes the interactions between p53, Mdm2, and DNA damage. An external stress induces a damage to the DNA, which promotes Mdm2 degradation. The level of p53 can then increase and activate DNA repair mechanisms. In parallel, p53 inhibits Mdm2 translocation from the cytoplasm to the nucleus through an indirect inactivation of AKT. However, in the nucleus, high level of p53 activates Mdm2 transcription, while Mdm2 induces the degradation of p53, thereby forming a negative feedback circuit. This figure has been drawn according to the Systems Biology Graphical Notation (SBGN) specifications.

## 3 Construction and analysis of the model

In this section, referring to the p53-Mdm2 network defined above, we introduce the different steps required for the definition of a logical model and for the analysis of its dynamical properties with the software GINsim, release 3.0.

### 3.1 GINsim

The GINsim software supports the definition, the simulation and the analysis of regulatory graphs, based on the (multilevel) logical formalism. GINsim is freely available for academic users from the GINsim website (http://ginsim.org), along with documentation and a model repository. For this tutorial, we use the recently released GINsim-3.0, which is available for all platforms with version 8 of the Java Virtual Machine. To get started with GINsim, download the corresponding Java ARchive (JAR file), with dependencies included, from the download section of GINsim website (http://ginsim.org/downloads). On your computer, double-click on the file icon to start the application or launch it with the command: java-jar GINsim-*#*version.jar in a terminal. Detailed instructions, troubleshooting and further options are available on the website.

### 3.2 Definition of a logical regulatory graph

Upon launch, GINsim displays a window enabling the creation of a new model, the import of a model in a supported format, or the opening of a previously defined model (if any). By clicking on the *New model* button, a window enabling the edition of a new logical regulatory graph opens.

To edit a graph, use the toolbox located just on the top of the window (below the menu bar). Passing slowly with the mouse on each of the editing tools pops a message explaining the function of each tool. Clicking on the *E* icon enables further edition of an existing node or arc upon selection, while the garbage can icon serves to delete selected arcs and nodes. Clicking once on one of the remaining icons enables the drawing of a single node or arc. Clicking twice on one of these tools will lock this mode, to draw several nodes or arcs without clicking repeatedly on the relevant tool.

#### 3.2.1 Definition of the regulatory nodes

First, we need to define nodes for the four key regulatory factors of the model: p53, Mdm2cyt, Mdm2nuc, and DNA damage (DNAdam). Each node has a unique identifier and a maximal level, defining a range of possible functional qualitative levels, as specified in Table 1. To define all the nodes in a row, first double-click on the node addition tool (symbol is a square with a plus sign) to lock this mode, then click four times on the panel to create the four nodes, with default identifiers and a maximal level of 1. Next, click on the *E* icon to stop adding nodes, and select each node to change its name and maximal level in the bottom edition panel. Figure 2 illustrates this step.

**Table 1:** Regulatory nodes and maximal levels for the p53-Mdm2 model.

| Regulatory nodes | Maximal levels |
| --- | --- |
| p53 | 2 |
| Mdm2cyt | 2 |
| Mdm2nuc | 1 |
| DNAdam | 1 |

**Figure 2:**
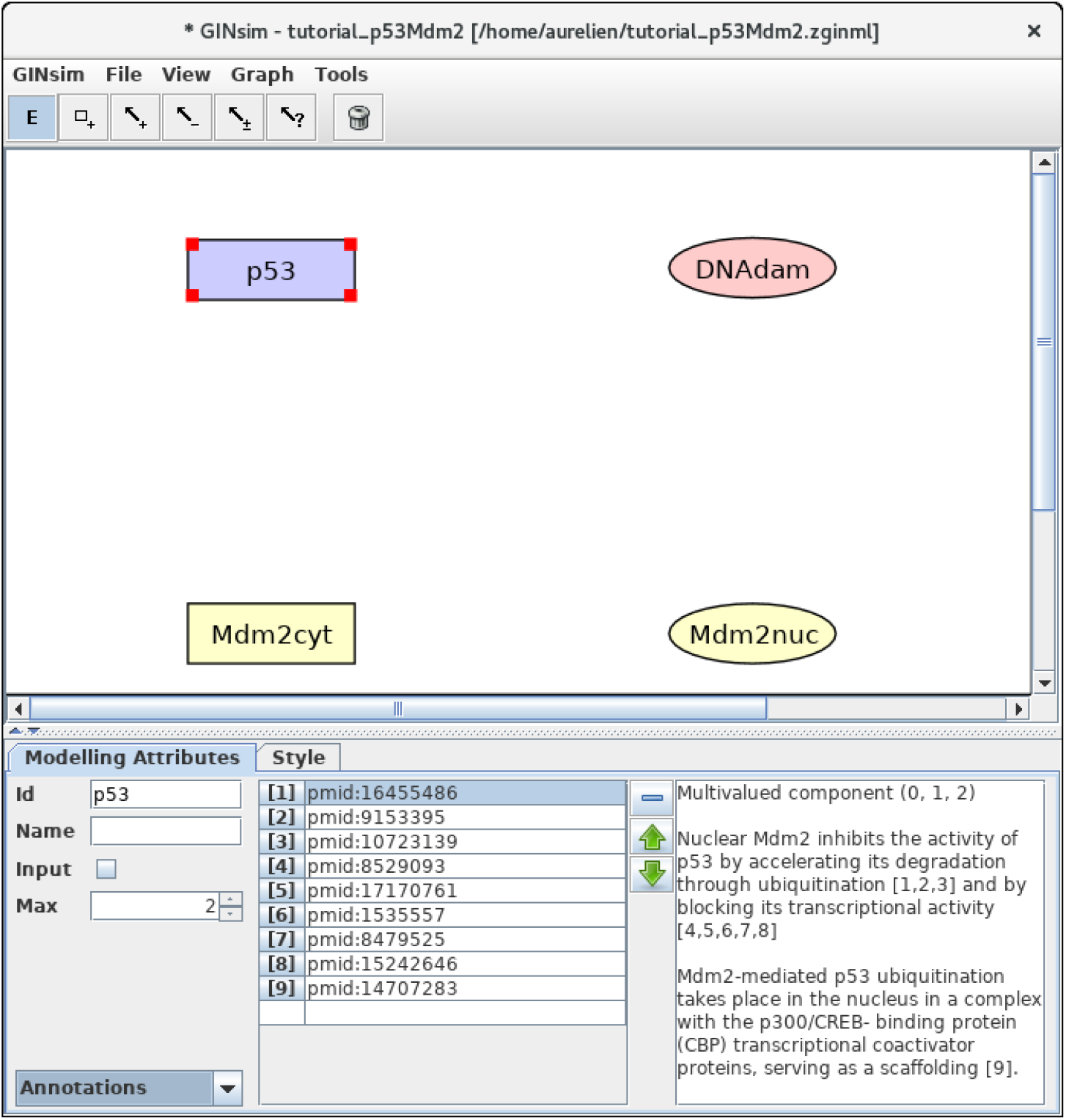
GINsim main window displaying the nodes of the p53-Mdm2 logical regulatory graph. The upper part of the window displays five scrolling menus. These menus provide access to classical file management options, as well as exports into various formats. The central area displays the regulatory graph (here the nodes of the p53-Mdm2 model), while the lower panel contains two tabs: the *Modelling Attributes* tab (selected here) and the *Style* tab, corresponding to the selected node, here p53. The graphical appearance of the nodes have been modified using the *Style* tab. The *Edit* button on the top is selected and emphasised with a green contour, enabling the edition of the attributes of the selected node, including its id and name, its maximal level (*Max*, here set to 2), and also the insertion of annotations in the form of free text (bottom right panel) or of links to relevant database entries (bottom middle panel).

#### 3.2.2 Definition of regulatory interactions

Next, we need to define the arcs representing the regulatory interactions between the factors considered in the model. An arc is defined by its source and target nodes, a sign, and a threshold, as described in Table 2 and illustrated in Figure 3. In the non-Boolean case, a node may have distinct actions on a target node, depending on its activity level (*e.g.*, from Mdm2cyt onto Mdm2nuc). In this case, one arc is drawn, which encompasses multiple interactions, each with its own threshold. An interaction is active when the level of its source is equal or above its threshold, but below the threshold of the next interaction. Add each arc between each relevant pair of nodes by selecting the relevant tool (addition of positive, negative, dual, or unknown interaction) and dragging a line from the source to the target node. Next, use the edition panel to specify their multiplicities and thresholds, an possibly change their signs.

**Table 2:** Interactions and thresholds for the p53-Mdm2 model.

| Source nodes | Target nodes | Thresholds | Interaction signs |
| --- | --- | --- | --- |
| p53 | Mdm2nuc | 1 | - |
|  | Mdm2cyt | 2 | + |
|  | DNAdam | 2 | - |
| Mdm2cyt | Mdm2nuc | 1 | + |
|  |  | 2 | + |
| Mdm2nuc | p53 | 1 | - |
| DNAdam | DNAdam | 1 | + |
|  | Mdm2nuc | 1 | - |

**Figure 3:**
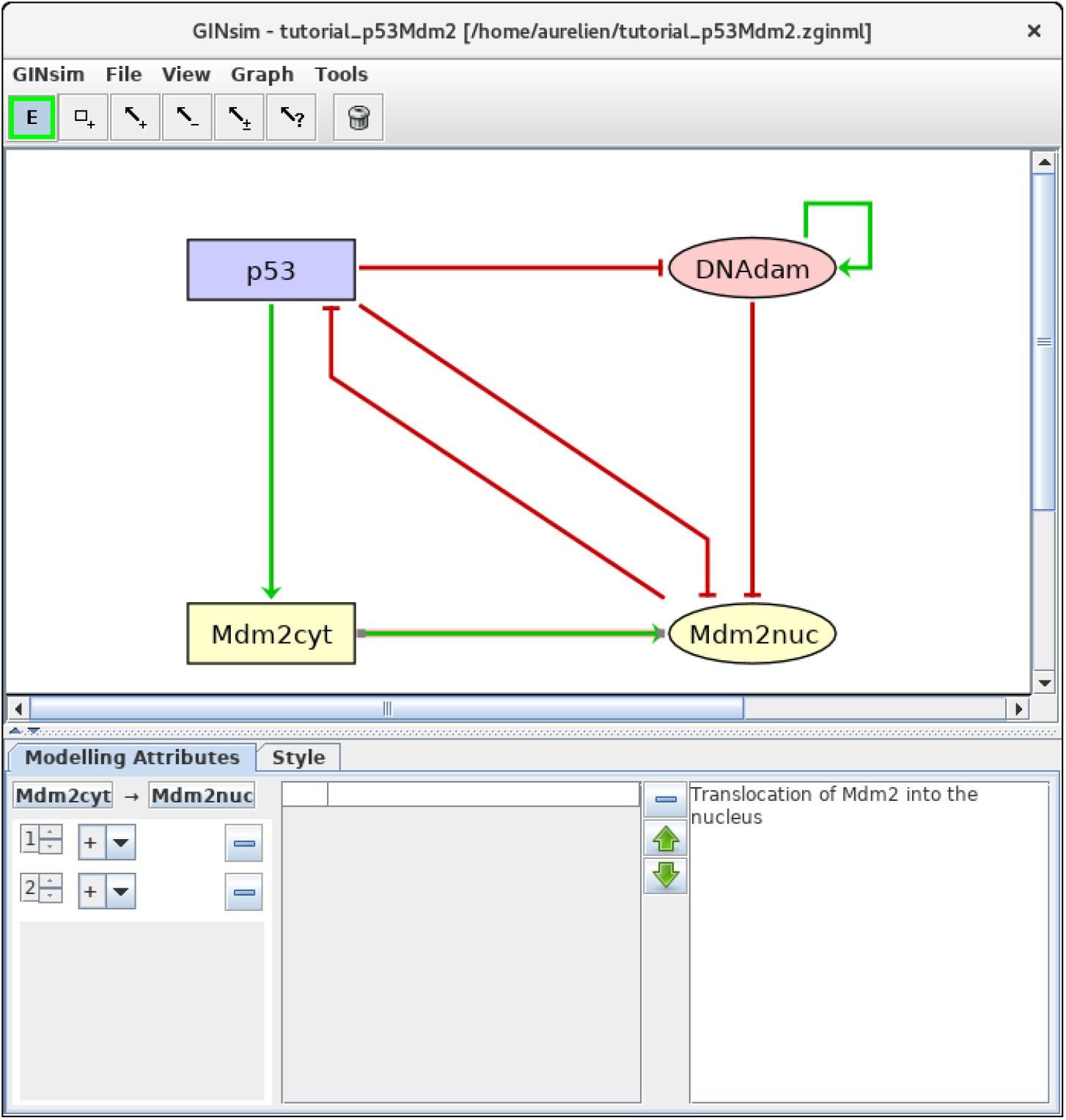
Regulatory arc management in GINsim. To add an arc, the corresponding arc button must be pushed (push twice to add several arcs in one go), allowing the drawing of an arc between a source node and its target. Once an arc has been defined, it can be further edited by selecting it along after locking the *E* button. The sign and threshold of the interaction(s) associated with an arc are defined within the *Modelling Attributes* tab, as shown here for the arc from Mdm2cyt onto Mdm2nuc. Note that an additional interaction of threshold level 2 can be created by clicking on the *+* button in the bottom left panel.

#### 3.2.3 Definition of the regulatory rules

We can now define the rules governing the evolution of the regulatory node levels. For each node, specify the logical rules listed in Table 3. For this, you need to select a node and select the *Formulae* view in the drop-down list at the bottom left of the GINsim window. Click on the little arrow in the main bottom panel, expand the tree view and then click on the *E* button, to enter a formula. Figure 4 illustrates this step. Note that the definition of adequate logical rules (or parameters, see see Note 1) is necessary to ensure the desired effects of each interaction on the target nodes. Per default, GINsim assigns a target value to each node devoid of explicit rule.

**Table 3:**
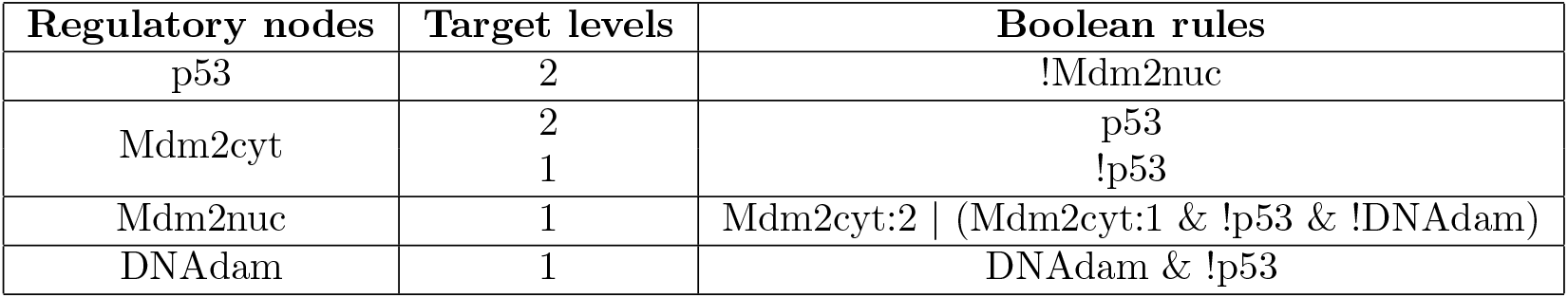
Logical rules for the nodes of the p53-Mdm2 model. This table lists the conditions enabling the activation of each node (up to level one in the case of a Boolean node, potentially up to higher levels for multilevel nodes, as for p53 and Mdm2cyt here). These conditions are defined in terms of Boolean expression using the NOT, AND and (inclusive) OR Boolean operators (denoted by !, & and | in GINsim, respectively).

**Figure 4:**
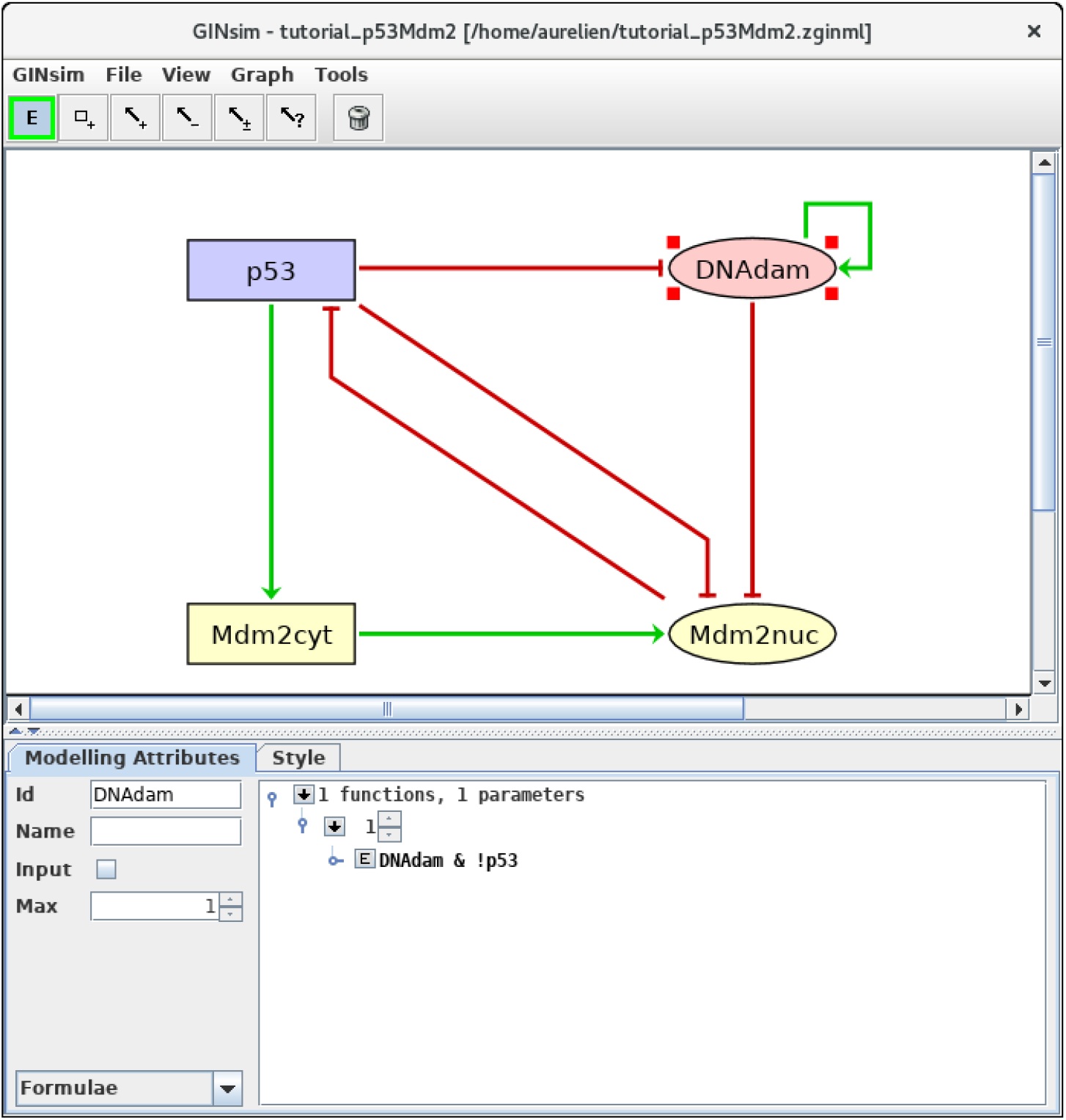
Defining logical rules for the regulatory nodes. This screenshot shows the *Modelling Attributes* associated with the selected node DNAdam. The maximal level is set to 1. After selecting *Formulae* with the bottom-left scrolling menu, the user can enter logical formulae after clicking on the little arrows in the main bottom panel. The target level (here set to 1 per default) can be changed in the case of a multilevel node. By clicking on the *E* button, one can directly write a formula, using literals (these should exactly match the IDs of nodes regulating the selected node, *i.e.* p53 or DNAdam in the present case) and the Boolean operators !, & and|, denoting NOT, AND and (inclusive) OR, respectively (following the usual priority ordering; parentheses can be used to define complex formulae). Note that several rows can be used in association with a single target value, while these rows are then combined with OR operators. Here, the formula *DNAdam* & !*p53* associated with the target value 1 implies that DNAdam will be maintained at a level 1 if already present, but only in the absence of p53.

#### 3.2.4 Adding annotations

To keep track of supporting data and modelling assumptions, you can add textual annotations and hyperlinks to relevant database entries, at the level of the model itself, as well as for each individual node or arc (see Figure 2 for an illustration). While the annotation panel is always visible when editing an arc, it requires to select the *Annotations* view (in the bottom left drop-down list) when editing a node.

#### 3.2.5 Changing layout and styles

You can change the layout and graphical appearance of nodes and arcs of the graph at your convenience. For this, select a node or an arc, along with the *Style* tab. You can either change the default style or define your own styles. To change the graph layout, drag a node to change its position or drag an arc to create a new intermediate point. An existing intermediate point can be moved or deleted using right-click.

#### 3.2.6 Node ordering

Selecting the Modelling Attributes tab, with no object selected in the main window, verify that the order of the nodes is: p53, Mdm2cyt, Mdm2nuc, DNAdam. If this is not the case, modify the node order accordingly, using the arrows close to the node list at the bottom-left of the tab. Using this node order will ease the comparison of your results with the Figures hereafter.

#### 3.2.7 Save your model!

The model along with simulations settings (see hereafter) can be saved into a compressed archive (with a zginml extension) by using the *Save* option in the *File* menu. Save your model regularly, as there is no undo functionality...

### 3.3 Dynamical analysis

The qualitative state of a logical model is defined by the activity levels of its nodes. At a given state, the rules associated with each node define its *target level*. When the current level of a node is different from its target level, it is called to update towards this target level, resulting in a transition to another state. Several nodes can be called for update at a given state.

Two main strategies are then commonly used. Under the *synchronous updating*, all concerned nodes change their levels simultaneously in a unique transition towards a single successor state. In contrast, the *asynchronous updating* generates a successor state for each node called for updating. If the current state involves *k* updating calls, it will thus have *k* successors, each differing from the current state by the level of a single node (see Note 2 for additional explanations). The introduction of priority classes allows to define subtler updating schedules (see Note 3 and Fauré et al. [16]).

The resulting state transitions define another type of graph called *state transition graph* (STG), which represents the dynamical behaviour of the logical model (*i.e.*, the regulatory graph + logical rules). In this graph, the nodes correspond to logical states, while the arcs represent state transitions induced by the rules along with the updating scheme. Using the default level layout of GINsim for transition graphs, it is easy to spot the stable states, defined as nodes with no outgoing arcs, displayed at the bottom. More complex attractors, defined as terminal *strongly connected components* (SCCs, maximal sets of nodes that are mutually reachable) denote oscillatory behaviours, which are harder to grasp visually.

Beyond the identification of attractors, we are particularly interested in knowing which of them can be reached from specific initial conditions. Such questions can be addressed by verifying the existence of *trajectories* (*i.e.*, sequences of transitions) from *e.g.*, initial to attractor states.

#### 3.3.1 Configuring a simulation

Selecting the *Run Simulation* option in the *Tools* menu opens a panel enabling the construction of the dynamics (see Figure 5).

**Figure 5:**
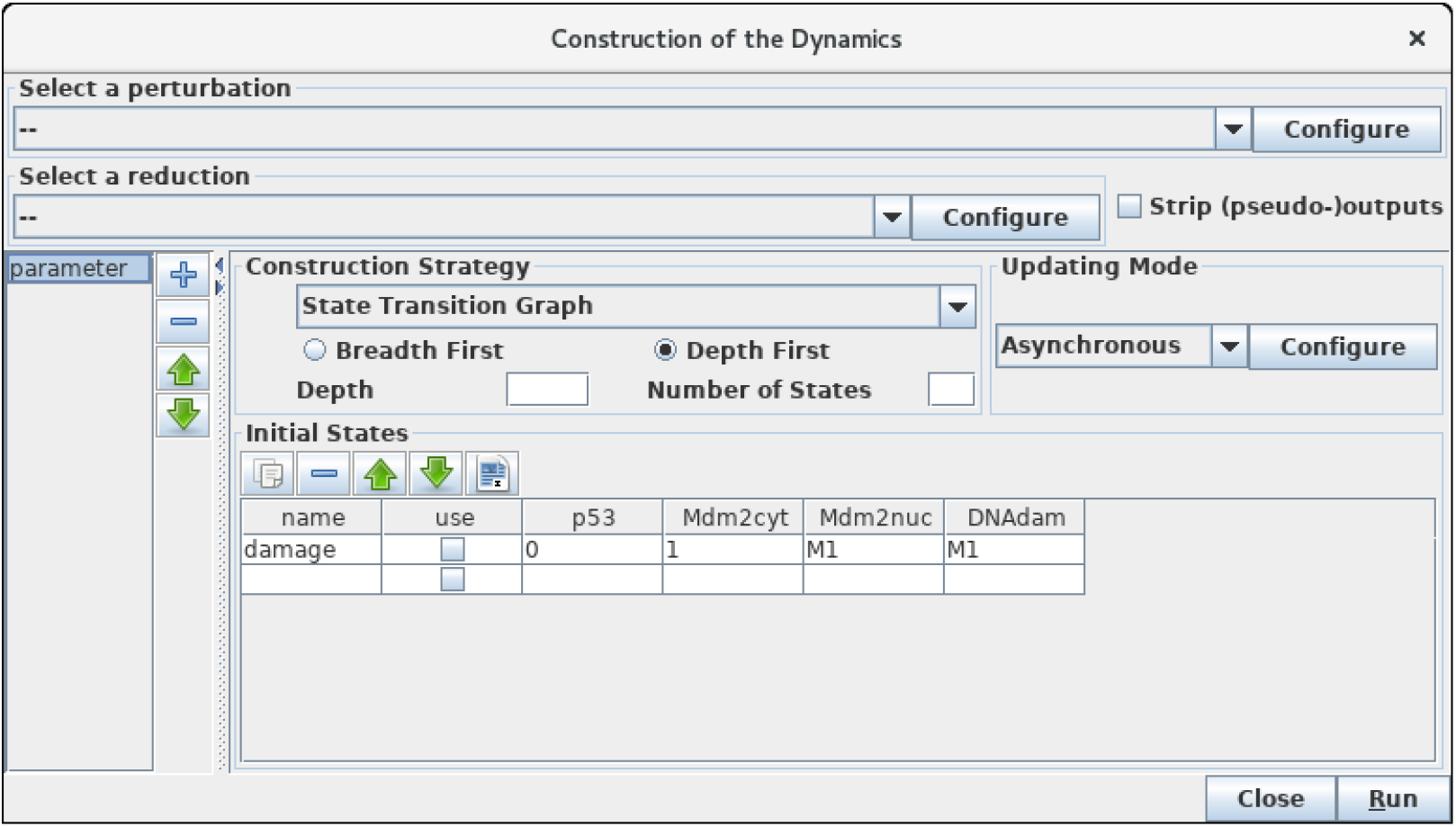
Launching of the construction of a state transition graph. This panel is obtained when selecting *Run Simulation* from the *Tools* scrolling menu in GINsim main window. The default simulation settings are shown, *i.e.* the construction of a state transition graph using the asynchronous updating, with no specified initial state (meaning that all states are considered in the simulation). Hitting the *Run* button will generate the corresponding state transition graph in a new window (see Figure 6). In the table under *Initial States*, one can define on or several initial states from which the dynamics will be constructed (just type the desired values in a row along with an optional name). Each row of the table defines a single (or a group) of states, and the check-boxes allow to select the states to be used for a simulation. The levels are specified for each node in the corresponding table cell. Nodes for which values are left free are denoted by stars (*). Initial states can be reordered, deleted and duplicated using the buttons just above the table. Here, a unique initial state has been defined, but not selected for simulation: the state 0111 (*i.e.* with p53 set to 0, and the three other nodes set to 1). Note that *M1* emphasises the fact that the value 1 is the maximal level for Mdm2nuc and for DNAdam. Several parameter configurations can be created and stored using the *+* button on the left side.

**Figure 6:**
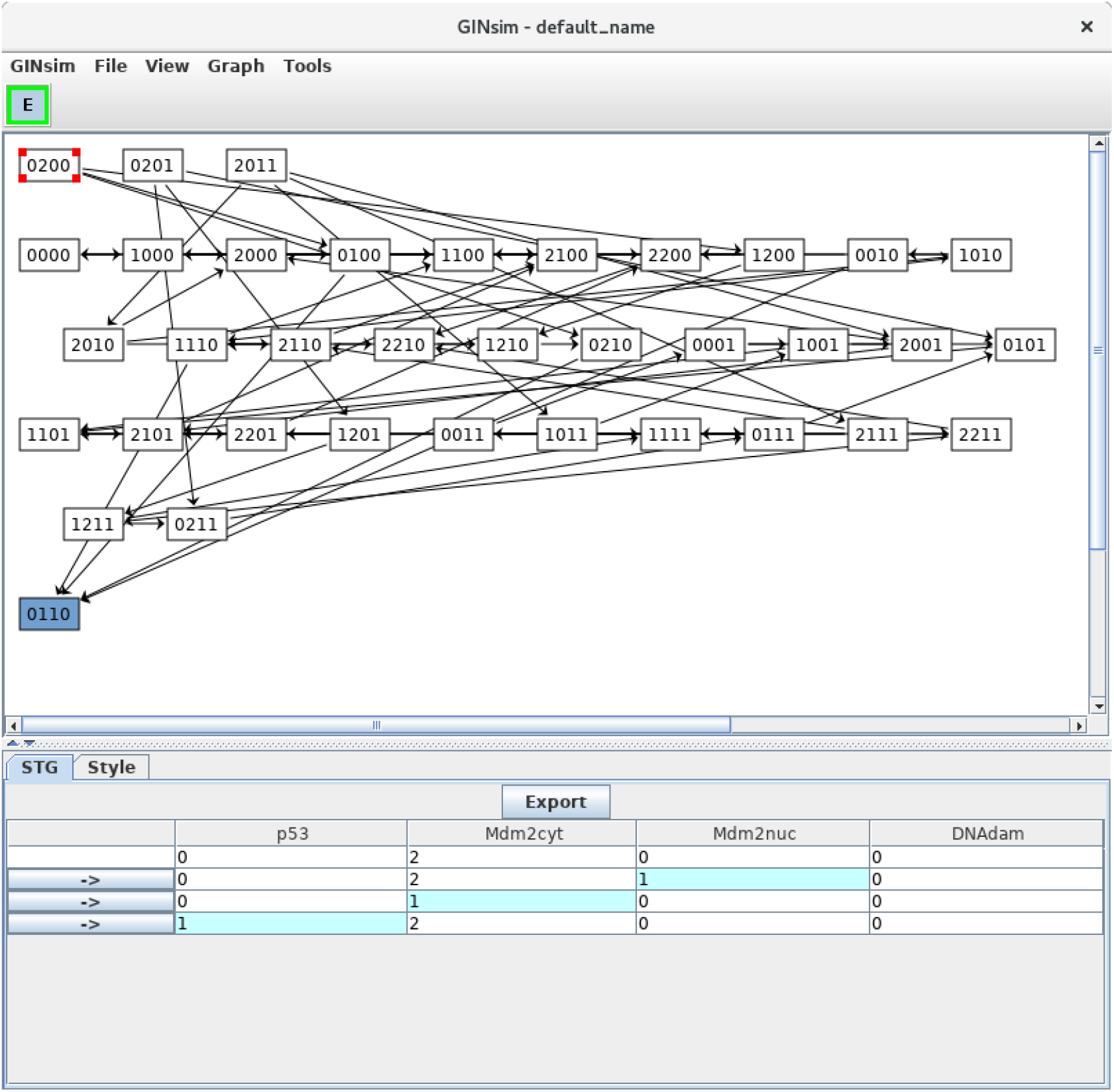
Asynchronous state transition graph for the p53-Mdm2 model. This STG has been generated with the simulation parameters shown in Figure 5, including the stable state 0110 laying at the bottom. The selected state 0200 is shown in the bottom panel, with its successors.

The boxes on the top of the panel labelled by *Select a perturbation* and *Select a reduction* permit to define (by clicking on the Configure buttons) and select (using the scrolling menus) model perturbations and reductions (see below).

The bottom left panel enables the definition and the recording of different parameter settings, which greatly facilitates the reproduction of results. One can create, delete and reorder parameter settings by using the buttons on the right of the panel.

Regarding the construction strategy, a scrolling menu enables the choice between the generation of a *state transition graph* (STG), its compression into a *strongly connected components graph* (SCC), or its further compression into a *hierarchical transition graph* (HTG) (for more details about these STG compressions, see [6]). Using another scrolling menu, the user can select the synchronous or asynchronous updating, or define or select predefined priority classes (see Note 3 for more details on priority classes).

Finally, the *Initial State* box on the bottom of the panel enables the definition and/or the selection of initial state(s), from which the construction of the dynamics will be performed. If no initial state is selected or specified, all the states will be considered in the simulation, leading to the construction of a full STG. As the number of possible states doubles with each additional (Boolean) node, the computation of the full STG is discouraged for models involving more than 15 nodes.

#### 3.3.2 Asynchronous simulations

Let us first consider the construction of the *asynchronous dynamics*. Before launching the simulation, check that the default settings are as specified in Figure 5: state transition graph, asynchronous updating, no perturbation selected, no initial state specified.

Clicking on the *Run* button launches the simulation, *i.e.* the computation of the state transition graph (STG). A dialog indicates that the result is available, allowing to display the STG or to perform other actions on it. For small STGs (up to 500 states), the default action opens a new window displaying the new STG with a simple layout. In the default level layout, the nodes with no incoming arc are placed at the top, whereas the nodes with no outgoing arc (*i.e.* stable states) are placed at the bottom. Stable states are further emphasised with a specific graphical attribute (by default an ellipse, whereas other states are displayed in rectangles). In this new window, you can re-arrange the nodes, either manually or by selecting a predefined layout in the View menu, change the graphical settings (clicking on the *Style* tab), and check outgoing transitions by selecting a state, as shown in Figure 6. Note that the scrolling menus propose various options, including path search functions, etc.

In Figure 6, the state 0200 (*i.e.*, with high level of Mdm2cyt, and the other three nodes OFF) is selected, from which three unitary transitions are enabled by the logical rules (Table 3): increase of Mdm2nuc from 0 to 1, decrease of Mdm2cyt from 2 to 1, and increase of p53 from 0 to 1. The selected state and its three successor states are shown in the bottom panel. It is possible to follow a transition path by clicking on a rightwards arrow button in the bottom panel, which switches the selection to the corresponding state. When the selected state also connects to predecessors states, these are also shown, preceded by leftwards arrows.

Note that a unique stable state was obtained, 0110 (following the order defined above, this vector states that p53=0, Mdm2cyt=1, Mdm2nuc=1 and DNAdam=0), which corresponds to the cell rest state (no p53, medium levels of cytoplasmic and nuclear Mdm2, no DNA damage).

#### 3.3.3 Direct computation of stable states

Select the *Compute stable states* option inn the *Tools* menu of the main window to verify that the unique stable state of this model is indeed 0110 (see Figure 7).

**Figure 7:**
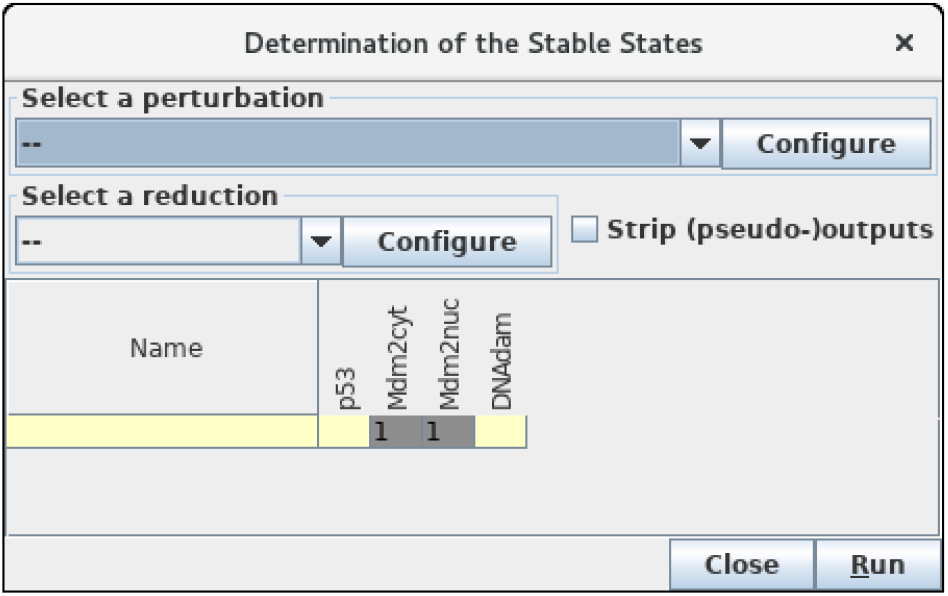
Determination of stable states. This window pops up upon selection of *Compute Stable States* with the *Tools* scrolling menu. After hitting the *Run* button, GINsim returns all stable states using an efficient algorithm. In the wild type case, we obtain a unique stable state 0110 as shown (yellow and gray cells denote levels 0 and 1, respectively).

This calculation uses an algorithm bypassing the construction of the STG, which is particularly useful for large models (see Naldi et al. [33] for more details).

If you get another (or no) stable state, check carefully the maximum level of each node, the threshold associated with each interaction, as well as each logical rule, as there must be a mistake somewhere...

#### 3.3.4 Synchronous simulations

For comparison, let us now build the state transition graph of the model using the synchronous updating strategy. Select *Run simulation* in the *Tools* menu of the main window, then select the *Synchronous* option with the scrolling menu under *Priority Class Selection* in Figure 5, and launch the simulation by clicking on the *Run button*.

The resulting STG (after a manual improvement of the layout) is shown in Figure 8. Naturally, the stable state 0110 is preserved (bottom left), but two cyclic attractors (bottom middle and right) are now obtained. Transitions representing single and multiple node updates are denoted by solid and dotted arcs, respectively.

**Figure 8:**
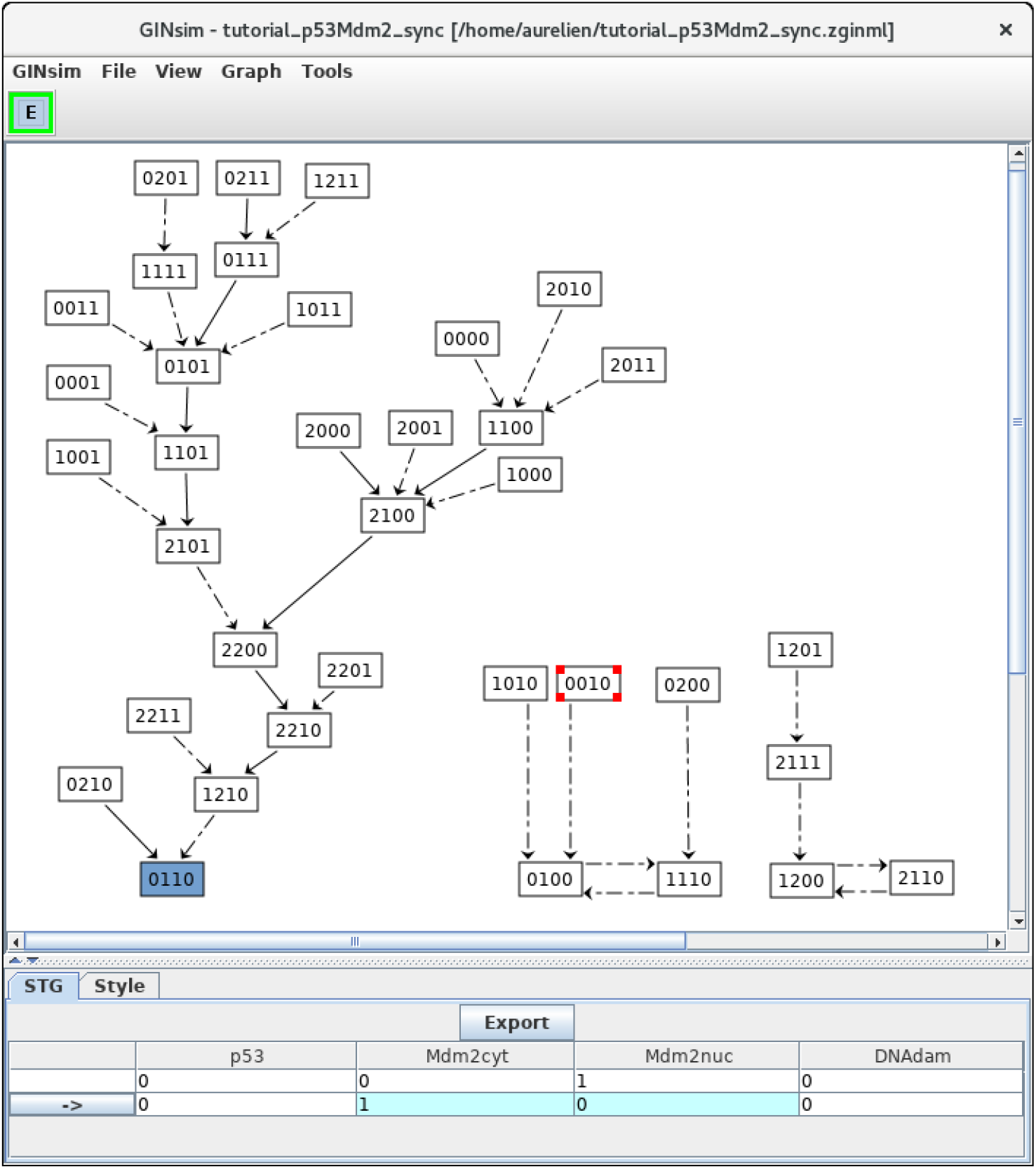
Synchronous state transition graph for the p53-Mdm2 model. This STG has been generated with the simulation parameters shown in Figure 5 (without specifying any initial state), but using the **synchronous** updating scheme). Note that the layout has been manually rearranged for sake of clarity. The STG is composed of three non connected subgraphs. On the left, we find back the resting stable state 0110, which can be reached from 26 other states. On the right, we see that the synchronous updating further generates two two-states cyclic attractors, which can be reached from three or two other states, respectively. Solid and dotted arrows denote single and multiple transitions, respectively.

Note that the selected state 0010 is leading to state 0100 through simultaneous changes of Mdm2cyt and Mdm2nuc, as shown in the bottom panel (blue cells).

#### 3.3.5 Compression of the STG

When the size of the model increases, the state transition graph (STG) quickly becomes hard to visualise. To ease its analysis, a compression (or compaction) can be performed by grouping sets of states into hyper-nodes. The arcs connecting the resulting nodes then still correspond to state transitions. In particular, by lumping states which belong to the same *strongly connected component* (SCC, in the graph-theoretical sense), an acyclic graph is obtained. Interestingly, the resulting SCC graph preserves the reachability properties of the original graph. However, in many situations, the SCC graph results only in a moderate STG compression.

To increase STG compression and ease the interpretation of the dynamics, we have recently introduced another acyclic graph, called *hierarchical transition graph*, which basically further merges linear chains of states (in addition to cycles) into single nodes [6]. The resulting graph preserves the attractors and other important dynamical properties, but do not fully conserve reachability properties.

Selecting the corresponding option with the Construction Strategy scrolling menu allows to compress the dynamics by using the hierarchical transition graph (HTG) representation. Figure 9 shows the resulting HTG, with all other simulation parameters maintained as shown in Figure 5.

**Figure 9:**
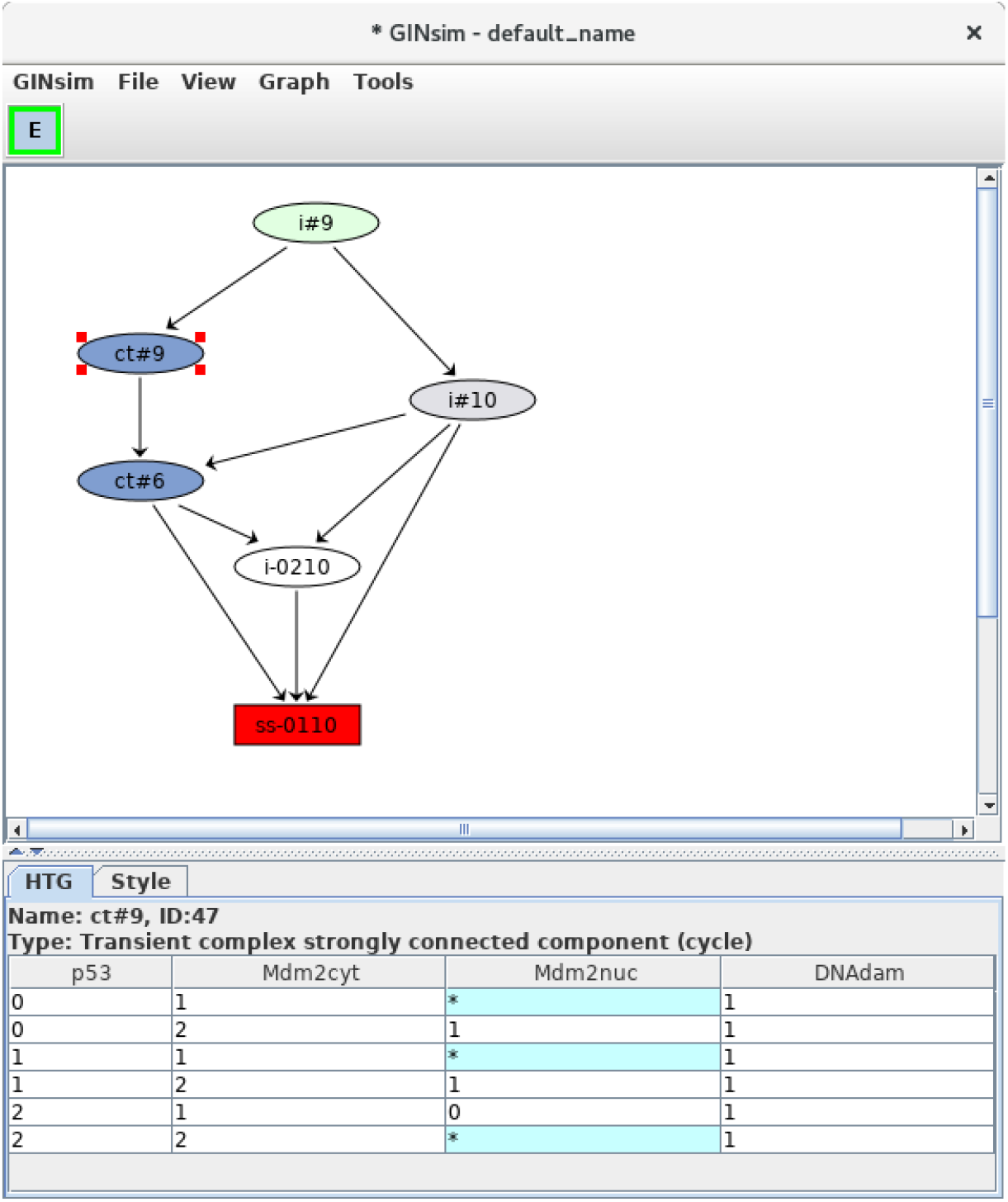
Hierarchical transition graph. The hierarchical transition graph for the complete asynchronous dynamics of the p53-Mdm2 model is shown. It has been obtained by selecting the construction of *Hierarchical Transition Graph* in the corresponding scrolling menu when launching the simulation. Note that the layout has been manually improved. The blue nodes correspond to the two non trivial strongly connected components of the STG, and the unique stable state is shown in red at the bottom. The blue node labelled by *ct#9* has been selected; this *transient cyclic component* encompasses nine states from the STG (as indicated by the *#9* in its name), which are listed in the bottom panel. The *\** denotes all possible values for the corresponding node. Hence the first row in the table listing the states encompassed by the hypernode *ct#9* corresponds to two states: 0101 and 0111.

Although relatively modest in this case (six nodes in the HTG, to be compared with 36 nodes for the original STG), this compression can be much more impressive in cases with long alternative trajectories (see *e.g.* Bérenguier et al. [6], Grieco et al. [23]). However, the computation of the HTG relies on that of the STG, with the compression done progressively. Hence, HTG computation may become intractable for large networks.

At the bottom of the HTG shown in Figure 9, note again the stable state 0110 (red box). In addition, two blue nodes representing strongly connected components can now be clearly seen, each labeled by *ct*, for *cyclic transient*, as both nodes are the sources of outgoing transitions.

The first of these cyclic components (ct#9) is selected and a list of the corresponding states is listed in the bottom panel (where a star stands for all possible values for the corresponding node, which enables a reduction of the list of states). This cyclic component contains nine states, all with the DNAdam node set to 1, p53 oscillating between the values 0 and 2, and both cytoplasmic and nuclear Mdm2 forms oscillating between the values 0 and 1. Hence, this cyclic component captures large oscillations of p53 in the presence of DNA damage.

The second cyclic component (ct#6) contains six states, with DNAdam now set to 0, with p53 and Mdm2cyt both oscillating between the values 1 and 2, and Mdm2nuc oscillating between the values 0 and 1. Hence, this cyclic component captures the smaller transient p53 oscillations observed just after DNA repair. In brief, starting from initial conditions with DNAdam=1, the system first goes through an unspecified number of large p53 activity oscillations, followed by DNA repair (DNAdam taking the value 0) along with transient smaller p53 oscillations, and finally the return to the rest state 0110.

### 3.4 Additional analyses

Several complementary analyses can be perfomed with GINsim. Hereafter, we illustrate three main functionalitie: the encoding of perturbations, an algorithm enabling the analysis of the roles of regulatory circuits, along with model reduction tool. Further information regarding GINsim functionalities can be found in the user manual and documentation available online.

#### 3.4.1 Definition of perturbations

Common perturbations are easily specified within the logical framework:

- A gene knock-down is specified by driving and constraining the level of the corresponding regulatory node to the value 0.
- Ectopic expression is specified by driving and constraining the level of the corresponding node to its highest value (or possibly to a range of values greater than zero, in the case of a multi-valued node).
- Multiple perturbations can be defined by combining several such constraints.
- More subtle perturbations can be defined by more sophisticate rewriting of node rules (*i.e.*, to change the effect of a given regulatory arc).

Various perturbations can thus be defined to account for experimental observations or to generate predictions regarding the dynamical role of regulatory factors or interactions. One can define perturbations to study the role of specific nodes or arcs in the dynamics of the model.

Define a mutant corresponding to an ectopic expression of DNAdam (see Figure 10). Such a perturbation can be encoded before the computation of stable states or of a state transition graph. Verify that the resting stable state 0110 is not stable anymore for this perturbation. Note the striking change of attractor for this perturbation, which now correspond to ample oscillations of p53, along with oscillations of both nuclear and cytoplasmic Mdm2 forms in the presence of DNA damage.

**Figure 10:**
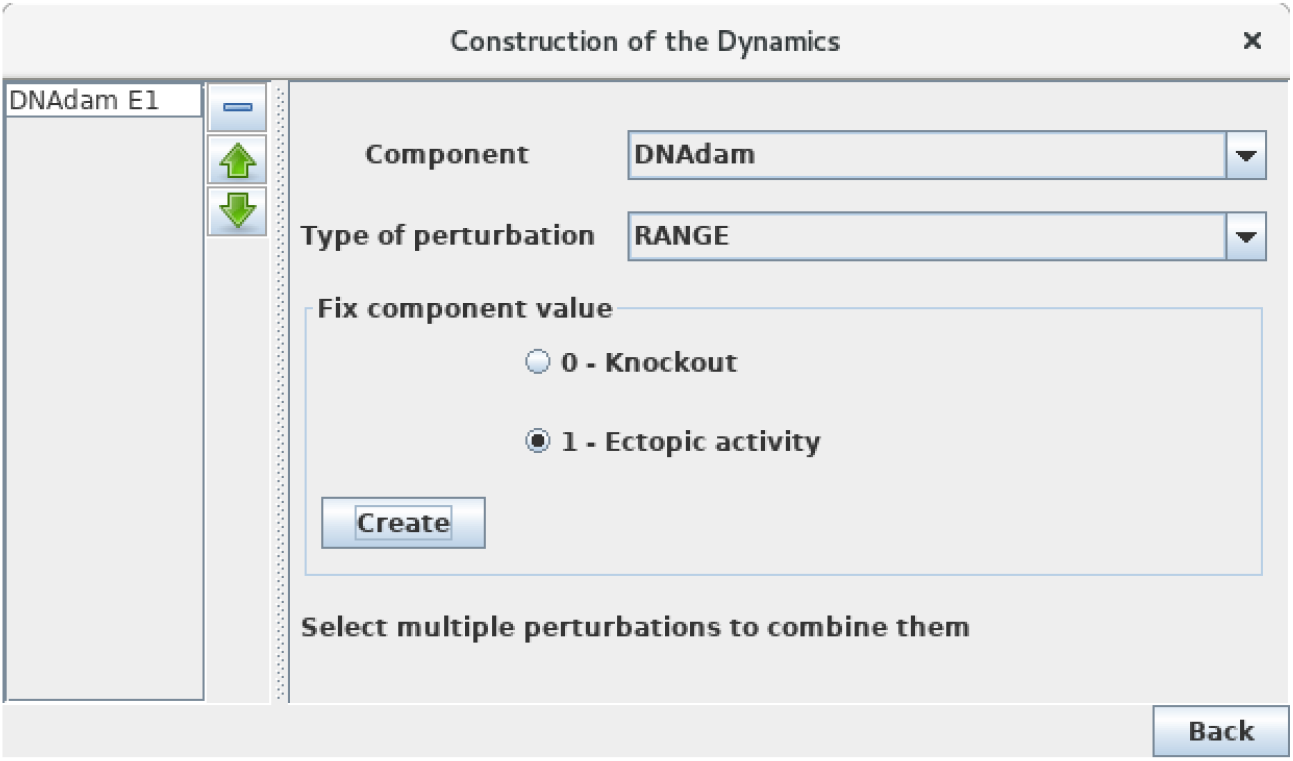
Perturbation specification. This window can be activated from the simulation launching window (Figure 5) and various other windows, including the *Compute Stable States* window. It enables the specification of various kinds of model perturbations, including loss-of-function and gain-of-function mutants. The figure illustrates the specification of a simple blockade of the level of DNAdam to level 1.

#### 3.4.2 Regulatory circuit analysis

Regulatory circuits are responsible for the emergence of dynamical properties, such as multistationarity or sustained oscillations (see Note 4). In this respect, GINsim implements specific algorithms to:

- Identify all the circuits of a regulatory graph (possibly considering constraints such as maximum length, consideration or exclusion of some nodes, etc.).
- Determine the functionality contexts of these circuits, using a computational method presented in Naldi et al. [33].

One can further identify and analyse the circuits of the model regulatory graph (see Subsection 3.2). Select the *Analyse Circuits* option of the *Tools* scrolling menu in the main window, then click on the *Search Circuits* button. Verify that the regulatory graph contains four circuits, among which three are functional (*i.e.*, have a non-empty functionality context). For each functional circuit, one can verify its sign and functionality context (depending on the rules), by clicking on the *Functionality Analysis* button. As shown in the Figure 11, the positive circuit defined by the cross inhibitions between p53 and Mdm2nuc is functional when Mdm2cyt=1 and DNAdam=0. Indeed, the inhibition of Mdm2nuc by p53 is not functional in the presence of DNAdam or of a high level of Mdm2cyt, or in the absence of Mdm2cyt.

**Figure 11:**
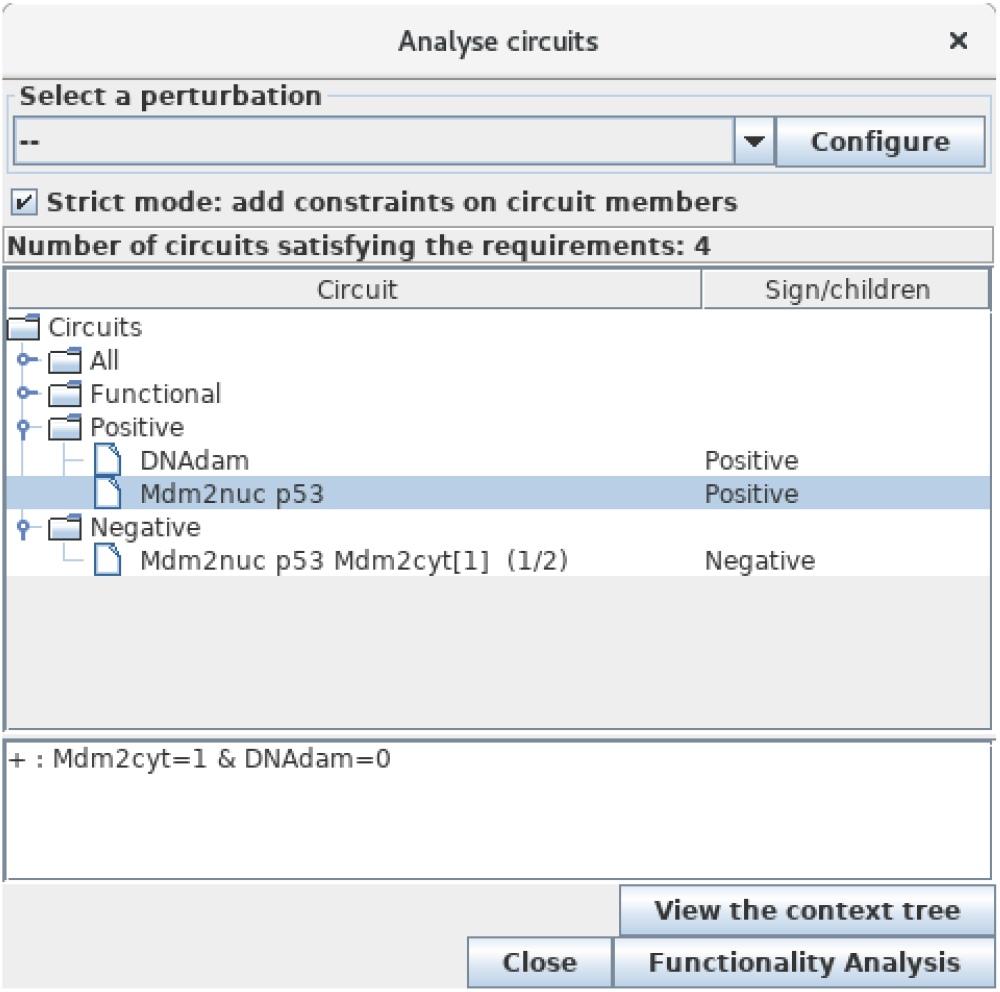
Circuit analysis for the p53-Mdm2 logical model. This window pops up after first selecting the *Analyse Circuits* option of the *Tools* scrolling menu in the main window, then clicking on the *Search Circuits* button, and finally launching the *Functionality Analysis* option. Among the four circuits found in the regulatory graph, three are functional: one is negative, while the other two are positive. The selected circuit (involving p53 and Mdm2nuc) is positive and functional when the level of Mdm2cyt is medium (equal to 1) in the absence of DNA damage (DNAdam=0).

#### 3.4.3 Reduction of logical models

When models increase in size, it quickly becomes difficult to cope with the size of the corresponding STG. One solution consists in simplifying or reducing the model before simulation. In this respect, GINsim implements a method to reduce a model on the fly, *i.e.*, just before the simulation. The modeller can specify the nodes to be reduced, and the logical rules associated with their targets are then recomputed taking into account the (indirect) effects of their regulators. This construction of reduced models preserves crucial dynamical properties of the original model, including stable states and more complex attractors [32].

For large networks, it might be useful to reduce the model before performing a simulation or other kind of dynamical analysis. Although our application is of limited size, we can still illustrate the use of GINsim model reduction functionality. Selecting the *Reduce Model* option in the *Tools* scrolling menu launches the reduction interface. Click on the *+* icon to define a reduction, then select the node Mdm2cyt for reduction, as shown in Figure 12. Clicking on the *Run* button generates a logical model encompassing only the three remaining nodes, where Mdm2nuc is the target of a dual interaction from p53. The logical rule associated with Mdm2nuc is consistently modified to take into account the former indirect effect of p53 through Mdm2cyt.

**Figure 12:**
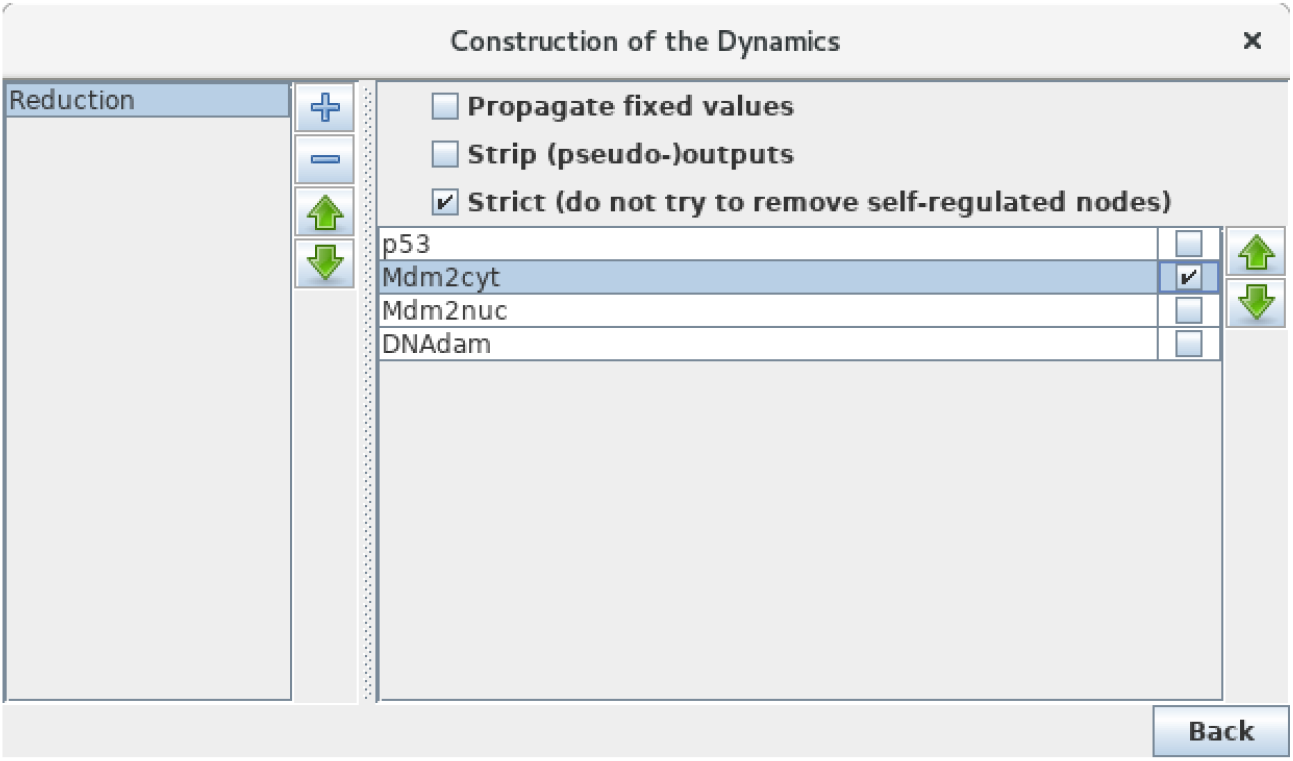
Model reduction. This window pops up following the selection of Reduce model from the Tools scrolling menu in the main GINsim window. Here, only Mdm2cyt has been selected for reduction. By hitting the *Run* Button, a reduced model is generated, provided that no self-regulated node is affected. Alternatively, one can close the window after the definition of one or several reduction(s) (the *+* button on the left enable to create new reductions) and then select a predefined reduction directly when performing simulations or other kinds of analyses.

Now that a reduction has been defined, you can select it when launching a simulation or computing stable states, without generating the reduced graph. Perform a complete asynchronous simulation to get the full state transition graph and verify that the number of states is now lower by a factor of three (12 states instead of 36) compared to Figure 6. You can further compute the HTG keeping the same parameter settings (asynchronous updating and full state space as initial condition). Although very much compressed, the resulting STG still captures the two kinds of transient oscillatory behaviour, ample in presence of DNA damage, smaller after DNA repair.

## 4 Conclusions

The logical formalism is particularly useful to model regulatory networks for which precise quantitative information is barely available, or yet to have a first glance of the dynamical properties of a complex model. For this protocol, we have considered a network comprising four regulatory factors, and we have followed the different steps enabling the delineation of a consistent logical model. Despite its limited size, this model yields relatively complex dynamics, including several transient oscillatory patterns and a stable state. It further served as a reference to illustrate advanced functions, such as model reduction or regulatory circuit analysis.

Large signalling networks have been handled with GINsim (*e.g.* [1, 8, 30]), in which input nodes denote external signals, which are not regulated and often maintained constant. Such *Input* nodes can be specified as such in GINsim to enforce the maintenance of the levels specified at initial states. As model reduction of input and output nodes or cascades have marginal impact on the dynamical properties [3], their reductions are facilitated in GINsim.

Furthermore, a novel functionality *Assess Attractor Reachability* in the Tool menu enables to evaluate the reachability of attractors based on stochastic simulation algorithms (for more details, see Henriques *et al.* article in this special issue).

Taking advantage of the multiple export formats supported by GINsim, it is also possible to use complementary tools, including stochastic simulations software (*e.g.* MaBoSS, see Stoll et al. [42]), model checking tools (*e.g.* NuSMV, see Abou-Jaoudé et al. [1, 3], Traynard et al. [48], or yet various graph visualisation and analysis packages (see Note 5 for a list of export options).

As mentioned in the introduction, various logical models for different cellular processes have been proposed during the last decades, many of them available in the repository included along with GINsim on the dedicated website (http://ginsim.org). The interested reader can thus download the model of his choice and play with it, reproduce some of the results reported in the corresponding publication, or modify and extend it according to his own research aims.

## 5 Notes

1. For each node, each combination of incoming interactions defines a logical parameter. This includes the situation in the absence of any specific activation or inhibition, or *basal level*. As a large fraction of the parameters are usually set to zero, this is the default value in GINsim (*i.e.*, any parameter lacking an explicitly assigned value is set to 0).
2. Transitions between states of the state transition graphs amount to the update of one (in the asynchronous case) or several (in the synchronous case) regulatory nodes. In any case, the update (increase or decrease) of a regulatory node is unitary (current value +1 or *−*1). Obviously, this remark applies only for multi-valued nodes (for which the maximal level is greater than 1).
3. Priority classes allow to refine the updating schemes applied to construct the state transition graphs [16]. GINsim users can group nodes into different classes and assign a priority rank to each of them. In case of concurrent updating transitions (*i.e.*, calls for level changes for several regulatory nodes in the same state), GINsim updates the node(s) belonging to the class with the highest ranking. For each priority class, the user can further specify the desired updating assumption, which then determines the treatment of concurrent transition calls inside that class. When several classes have the same rank, concurrent transitions are treated under an asynchronous assumption (no priority).
4. A regulatory circuit is defined as a sequence of interactions forming a simple closed directed path. The sign of a circuit is given by the product of the signs of its interactions. Consequently, a circuit is positive if it has an even number of inhibitions, it is negative otherwise. R. Thomas proposed that positive circuits are necessary to generate multistationarity, whereas negative circuits are necessary to generate stable oscillations (see Thieffry [44] and references therein). External regulators might prevent the functioning of a circuit imbedded in a more complex network. Naldi et al. [33] presents a method to determine the *functionality context* of a circuit in terms of constraints on the levels of its external regulator. A circuit functionality context can be interpreted as the part of the state space where the circuit is functional, *i.e.*, generates the expected dynamical property [13].
5. GINsim allows the user to export logical regulatory graphs as well as state transition graphs towards various formats, facilitating the use of other software:
  - SBML-qual, the qualitative extension of the popular model exchange format [9].
  - MaBoSS, a C++ software for simulating continuous/discrete time Markov processes, applied on a Boolean networks (https://maboss.curie.fr/).
  - BoolSim (http://www.vital-it.ch/software/genYsis/).
  - GNA, a software for the piecewise linear modelling of regulatory networks (http://ibis.inrialpes.fr/article122.html).
  - NuSMV, a symbolic model-checking tool (http://nusmv.fbk.eu/).
  - Integrated Net Analyzer (INA) supporting the analysis of Place/Transition Nets (Petri Nets) and Coloured Petri nets (http://www2.informatik.hu-berlin.de/~starke/ina.html).
  - Snoopy, a tool to design and animate hierarchical graphs, among others Petri nets (http://www-dssz.informatik.tu-cottbus.de/DSSZ/Software/Snoopy).
  - Graphviz, an open source graph visualization software offering main graph layout programs (http://www.graphviz.org/).
  - Cytoscape, a popular open source software platform for visualizing molecular interaction networks (http://www.cytoscape.org/).
  - Scalable Vector Graphics (SVG) format, an XML standard for describing two-dimensional graphics (http://www.w3.org/Graphics/SVG/).

## Author Contributions

While AN and PTM have been the main developers of GINsim over the last years, all authors of this manuscript have taken part in various practical tutorials introducing the usage of GINsim to biologists, which served as a basis for the preparation of this method article. All authors have further participated in the writing of the manuscript and in the preparation of the Figures. AN, CH, WAJ and PTM should be considered as co-first authors, and CC and DT as co-last authors. All authors reviewed the content of this article and agreed to endorse it.

## Acknowledgments

The authors acknowledge numerous constructive comments from GINsim users over the years, in particular insightful feedback from Laurence Calzone, Samuel Collombet, Karla Corral, Adrien Fauré, Swann Floćhlay, Asmund Floback, Anna Niarakis, Elisabeth Remy, Otoniel Rodriguez, and Gautier Stoll.

## Supplemental Data

The p53-Mdm2 model can be found in GINsim model repository, along with annotations, at the url: http://ginsim.org/model/p53-Mdm2.

The stable release of GINsim 3.0 can be freely downloaded from http://ginsim.org/downloads, as a single jar file will all dependency libraries included.

